## Supplementary material for "Insights from Molecular Docking and Dynamics Simulations of P2RX7-αSyn Complex": Supplimentary figures

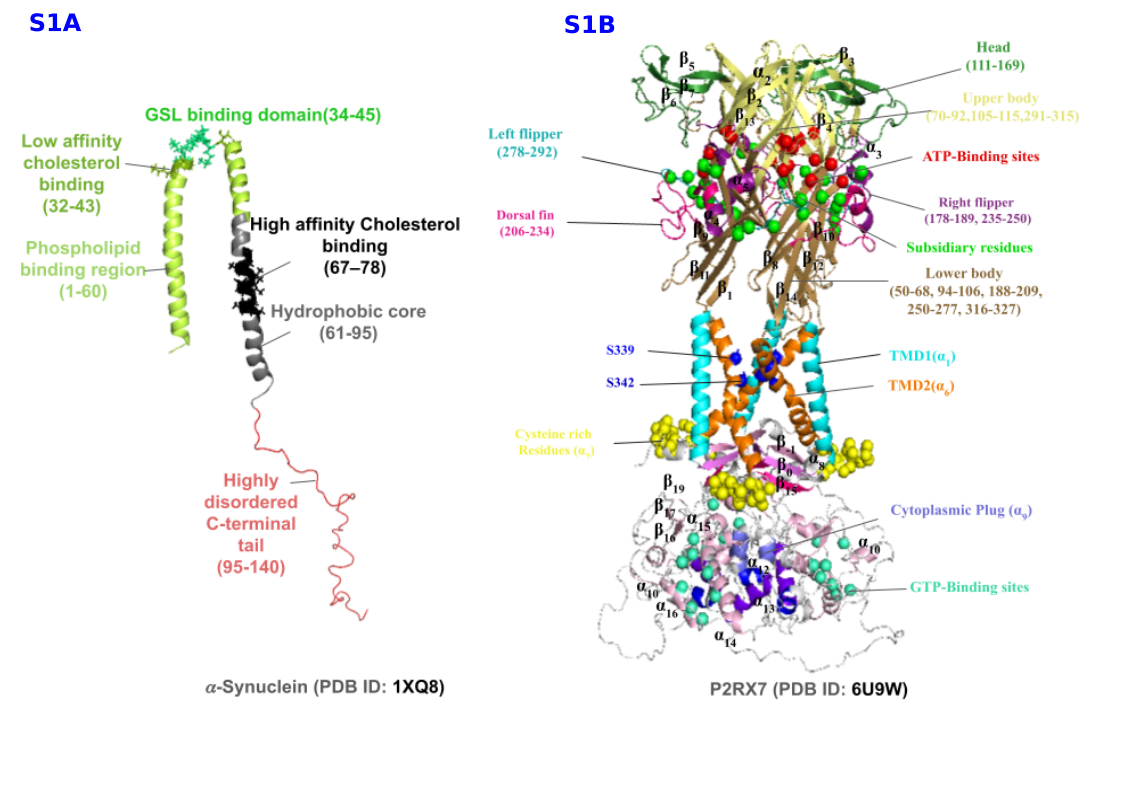

**Supplementary Figure 1:** (A) Structural features annotation of SNCA (PDB ID 1XQ8) and P2RX7 (PDB ID 6U9W).

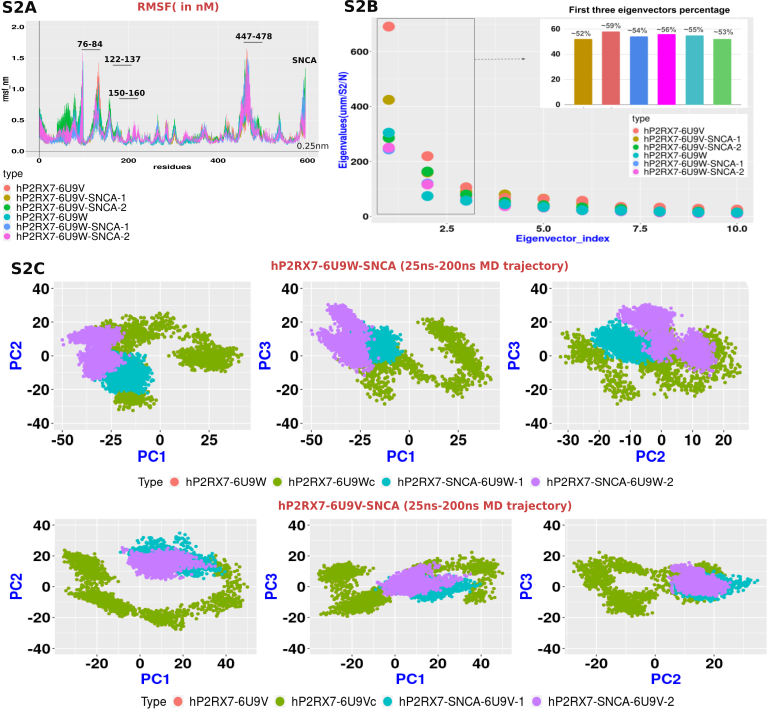

**Supplementary Fig.2 Dynamical changes induced by** 𝛼Syn **on P2RX7 in larger trajectory:** (A) RMSF of apoP2RX7 and P2RX7-SNCA complex models after 200 ns MD simulations. There are certain regions in P2RX7 in P2RX7 in both apo and complexes which are quite flexible. Most of these regions (Head, ATP binding sites, cytoplasmic ballast). (B) Distribution of first 10 eigenvectors of all the models after PCA analysis of 175 ns MD trajectory. First three eigenvectors of each model within the box show their proportion in the bar diagram labeled with eigenvector percentage. (C) Concatenated trajectory of respective apoP2RX7 and P2RX7-SNCA complexes of larger 175 ns MD simulations were used for PCA analysis. The projection of PC1 vs PC2, PC1 vs PC3, and PC2 vs PC3 for concatenated trajectories of apoP2RX7 and the P2RX7-αSyn (SNCA) complex using eigenvectors shows that certain subsets of conformations overlap in the apoP2RX7 and P2RX7-αSyn complexes, particularly in the central regions of the plots. This overlap suggests structural similarities between the apoP2RX7 and the P2RX7-αSyn complex. However, in the PC1 vs PC2 plot, isolated pockets are located away from the center, indicating structural dissimilarities. This observation implies that while there are common structural features, some regions of the conformational space exhibit notable differences between the Apo and bound states.

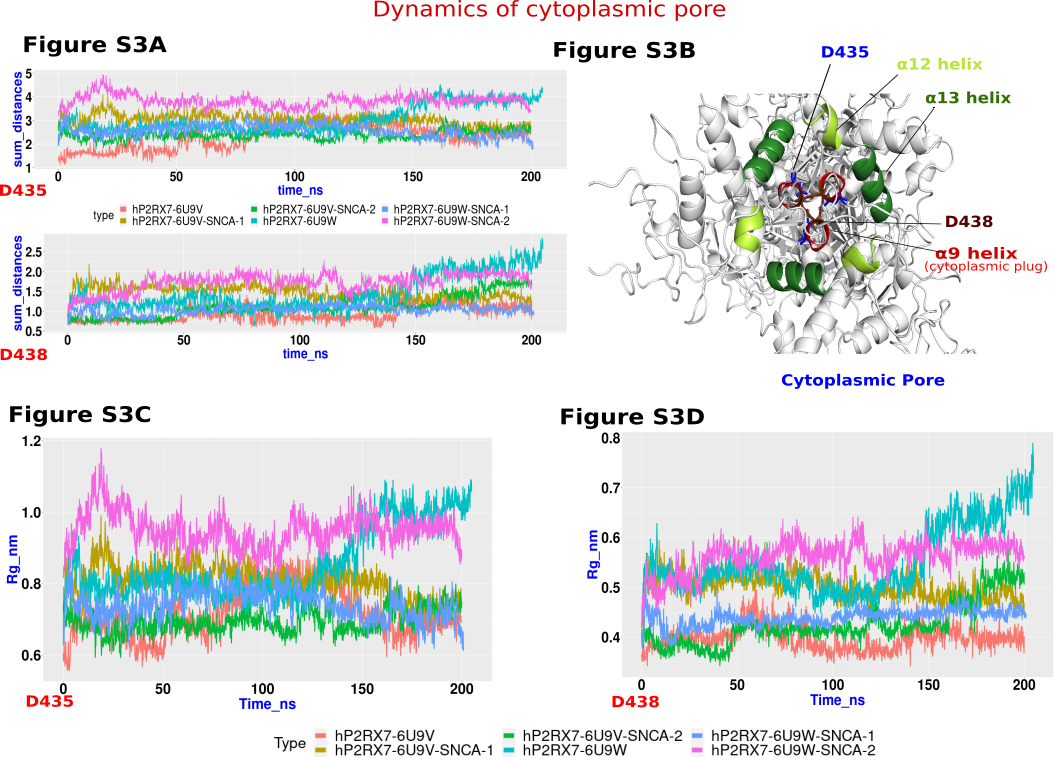

**Supplementary figure 3;** Cytoplasmic pore dynamics: A) Perform a pairwise distance analysis for residues D435 and D438 in chain A, comparing these distances to the corresponding residues in chains B and C within both the apo and P2RX7-SNCA complexes using GROMACS software. After computing the distances, sum them separately for D435 and D438 and visualize the results with a line plot. B) Illustrate the cytoplasmic pore structure, focusing on the α12- and α13-helices, and the cytoplasmic plug represented by the α9-helix, using cartoon diagrams. C) and D) Conduct radius of gyration (Rg) analysis for residues D435 and D438 in both the apo and P2RX7-SNCA complexes.

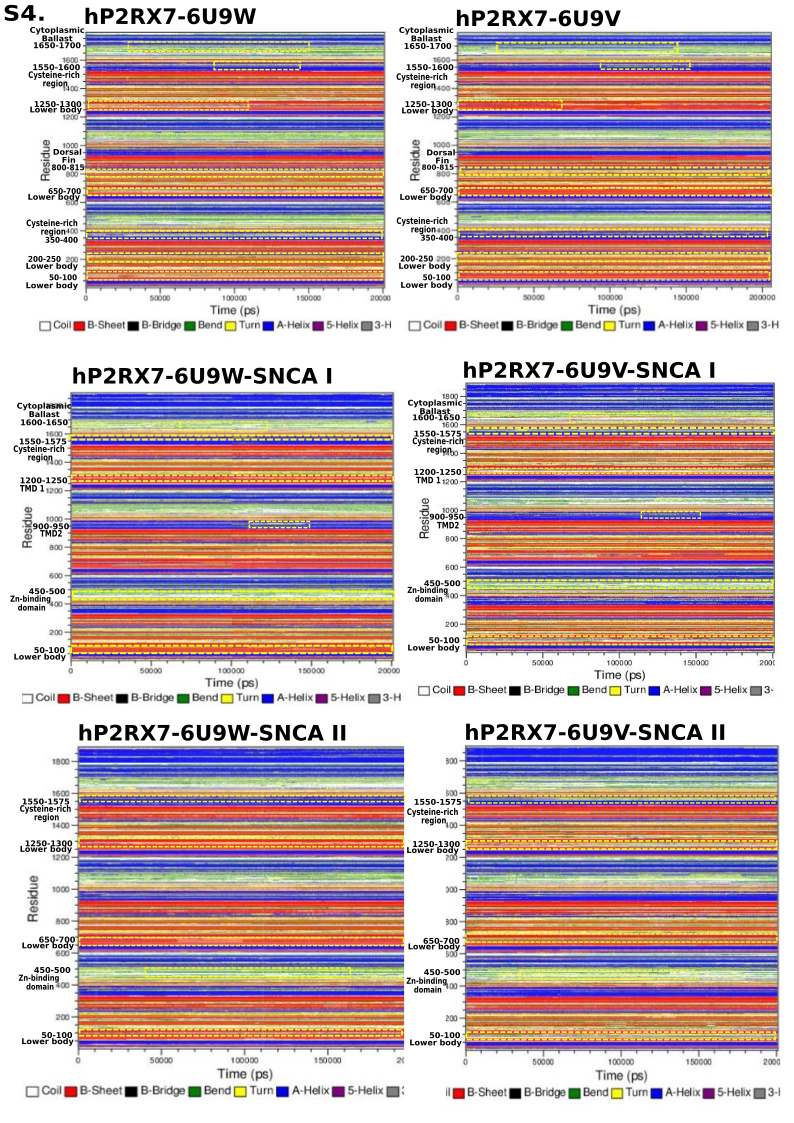

**Supplementary Fig.4 Secondary structures contents analysis.** (A) The overall changes in the distance between S339 and S342 of the respective chains in trimerized P2RX7 were calculated using the minimum distance analysis in GROMACS. The figure illustrates the distances between these residues across chains, with alterations during MD simulations quantified. No significant alterations in distance were observed throughout the simulations. (B) A cartoon diagram of the transmembrane domain (TMD) of P2RX7, formed by the α1 and α6 helices, highlights the distance between the TMD residues S339 and S342 across respective chains, which were subjected to pairwise distance analysis. (C) The α-helix content of α1 and α6 was quantified using the ALPHARMSD function in the PLUMED software, which estimates the total number of α-helix contents in the MD trajectory over specific time intervals. The analysis revealed a reduction in the α6 (TMD2) content, indicating unfolding of the TMD2 α-helix. (D) Similar results were observed in the quantification of α1 and α6 content using the DSSP program, confirming the reduction in TMD2 α-helix content. (E) The cytoplasmic cap, formed by the β-1, β0, and β15 strands, is critical for TMD pore opening. The AlphaBeta collective variables, which measure the phi and psi angles of specific secondary structures relative to standard angles in PLUMED, were used to assess the unfolding of beta strands. However, no significant changes were detected in this study. (F) A cartoon representation of the β-1, β0, and β15 strands forming the cytoplasmic cap is provided. (G) The cytoplasmic pore, formed by the α12 and α13 helices, and the cytoplasmic plug (α9) hanging over the pore, were analysed. The stability of these α-helices was measured using the ALPHARMSD, which quantified their contents throughout the entire MD trajectory for each model. The results were compared between apoP2RX7 and P2RX7-SNCA complexes, with no significant changes observed. (H) A cartoon representation of the cytoplasmic pore components (α12 and α13) and the cytoplasmic plug is shown.

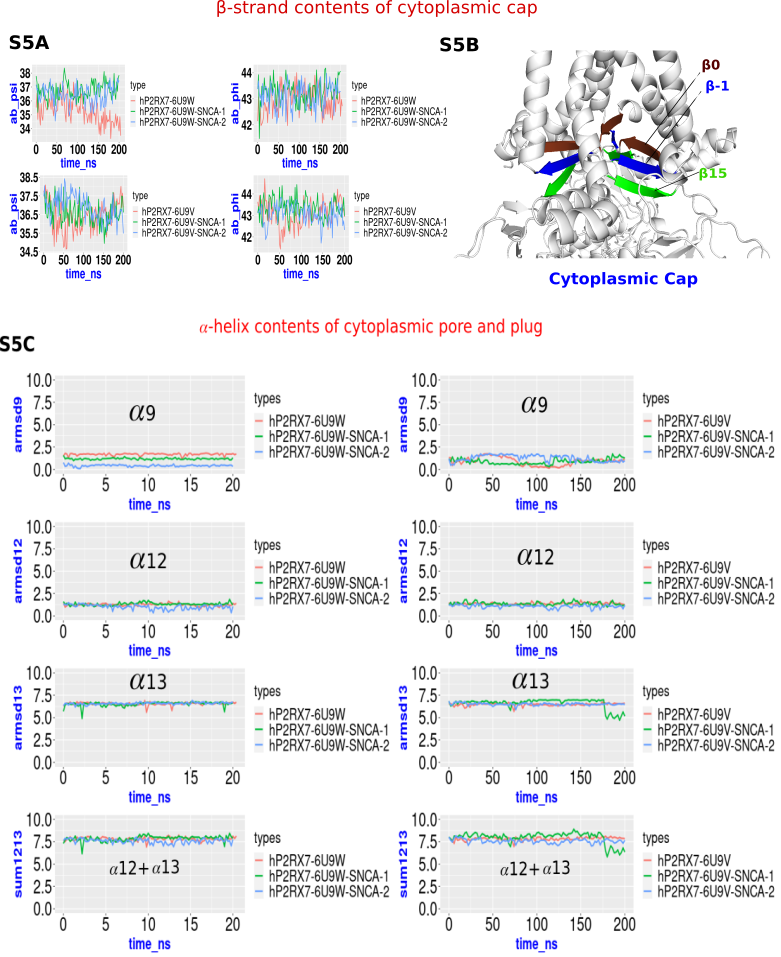

**Supplementary Figure S5:** The evaluation of cytoplasmic cap stability; A) The Alpha-Beta Collective variables from the PLUMED plugin were used to assess the alpha and beta strand contents by analyzing standard phi and psi angles of -135° and 135° in both the apo and P2RX7-SNCA complexes. The results are displayed as a line plot. The graph shows in upper panel, the open forms of apoP2RX7 in dark orange while the corresponding SNCA complexes are shown in dark olive green hP2RX7- 6U9W- SNCA-1) and cornflower blue (hP2RX7- 6U9W- SNCA-2). The same colour code was depicted in lower panel for apo forms of closed P2RX7 and corresponding complexes (hP2RX7- 6U9V1- SNCA-1 and hP2RX7- 6U9V1- SNCA-2). B) Cartoon diagrams illustrate the cytoplasmic cap, which is composed of β-1, β0, and β15 beta strands, colour blue, maroon, , and green, respectively. C) The α-helix content of the cytoplasmic pore (α12 and α13) and the cytoplasmic plug (α9) was calculated using αRMSD and presented as a line graph. The color code was same as shown in figure A.

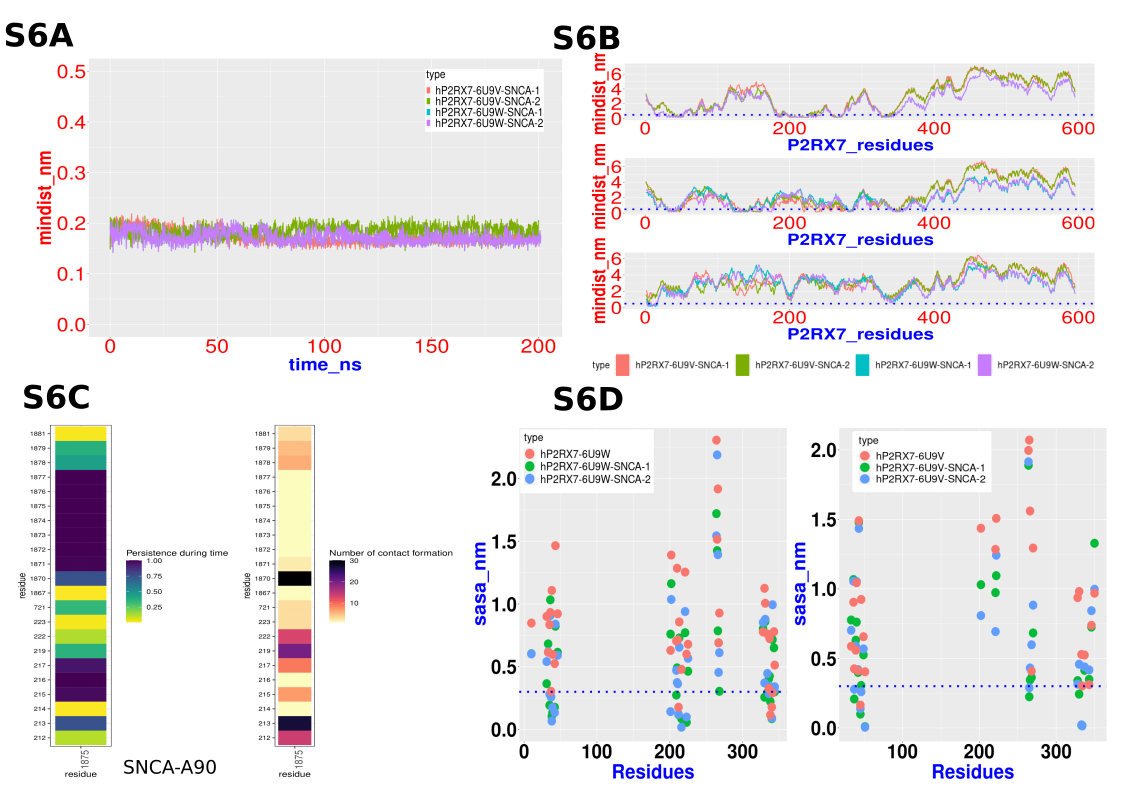

**Supplementary Figure S6:** The evaluation of contacts between P2RX7 and α-Syn; A) The minimum distance analysis between α-Syn and P2RX7 showing quite close proximity. The results are displayed as a line plot. The graph shows corresponding P2RX7-SNCA complexes are shown in dark orange (hP2RX7- 6U9V- SNCA-1), dark olive green (hP2RX7- 6U9V- SNCA-2), corn blue (hP2RX7- 6U9W- SNCA-1) and purple (hP2RX7- 6U9W- SNCA-2). B) Minimum distances of each residues of P2RX7 to the α-Syn shows certain region of P2RX7 quite close to the α-Syn, most of these regions are trans membrane domains in chain A and chain B. The same colour codes of legend were followed shown in previous figure. C) The contact analysis by CONAN using 5Å distance cut-off shows persistence of contacts and number of contact formation. D) Solvent Accessibility Surface Area (SASA) analysis of P2RX7 residues shows proximity to the α-Syn. Most of the residues show higher solvent accessibility**.** The graph in left panel shows the open forms of apoP2RX7 in dark orange while the corresponding SNCA complexes are shown in dark olive green hP2RX7- 6U9W- SNCA-1) and cornflower blue (hP2RX7- 6U9W- SNCA-2). The same colour code was depicted in right panel for apo forms of closed P2RX7 and corresponding complexes (hP2RX7- 6U9V1- SNCA-1 and hP2RX7- 6U9V1- SNCA-2).

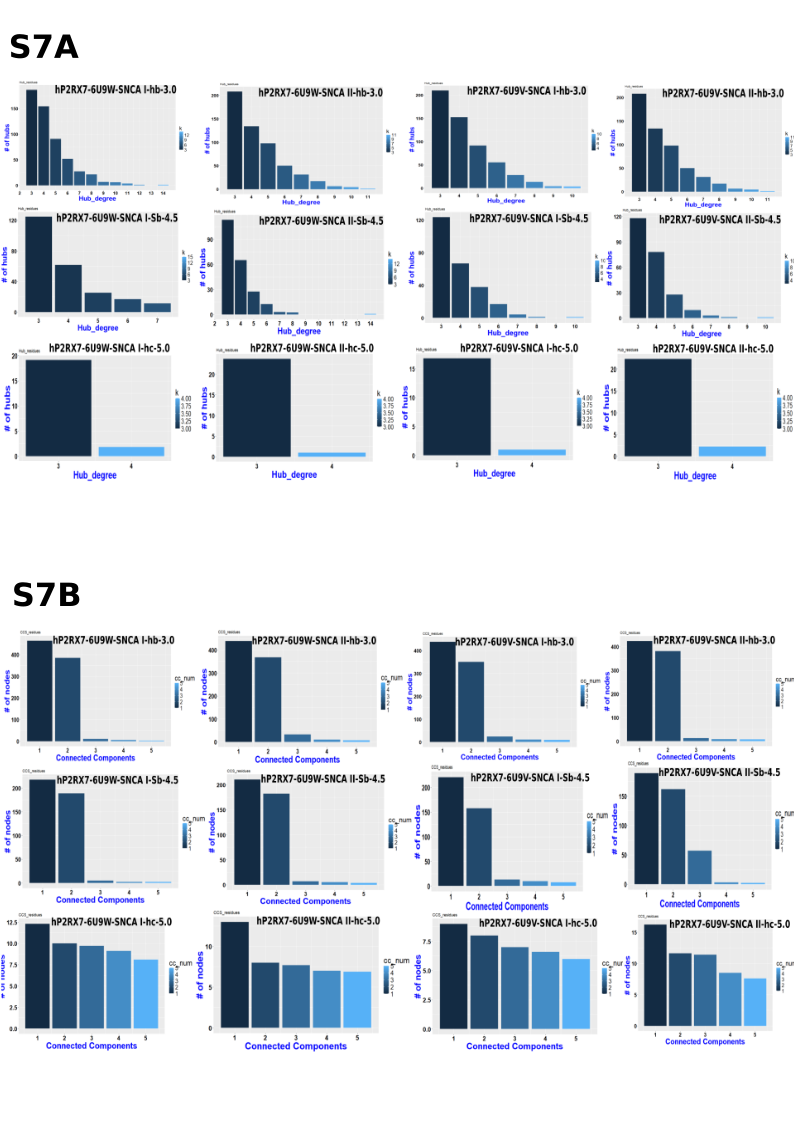

**Supplementary Figure7:** Hub and connected component analysis of P2RX7-SNCA Complex using Protein Structure Network (PSN) Analysis; (A) The number of hub residues identified from the PSN analysis of P2RX7-SNCA complexes, specifically hP2RX7-6U9W-SNCA and hP2RX7-6U9V-SNCA. The PSN was constructed using contact information derived from salt bridge interactions (cut-off 4.5 Å), hydrogen bonds (cut-off 3.5 Å), and hydrophobic contacts (cut-off 5 Å) obtained from MD trajectories. These hub residues are crucial for maintaining the structural integrity and function of the protein complex. (B) The top 5 connected components identified from the PSN analysis of the hP2RX7-6U9W-SNCA and hP2RX7-6U9V-SNCA complexes. These components represent the most interconnected and stable regions of the protein structure during the 200 ns MD simulations, indicating key regions of structural and functional significance within the complex.
